# Application of tissue-scale tension to avian epithelia *in vivo* to study multiscale mechanics and inter-germ layer coupling

**DOI:** 10.1101/2024.04.04.588089

**Authors:** Panagiotis Oikonomou, Lisa Calvary, Helena C. Cirne, Andreas E. Welch, John F. Durel, Olivia Powell, Kwantae Kim, Nandan L. Nerurkar

## Abstract

As cross-disciplinary approaches drawing from physics and mechanics have increasingly influenced our understanding of morphogenesis, the tools available to measure and perturb physical aspects of embryonic development have expanded as well. However, it remains a challenge to measure mechanical properties and apply exogenous tissue-scale forces in vivo, particularly for epithelia. Exploiting the size and accessibility of the developing chick embryo, here we describe a simple technique to quantitatively apply exogenous forces on the order of 1-100 N to the endodermal epithelium. To demonstrate the utility of this approach, we performed a series of proof-of-concept experiments that reveal fundamental and unexpected mechanical behaviors in the early chick embryo, including mechanotype heterogeneity among cells of the midgut endoderm, complex non-cell autonomous effects of actin disruption, and a high degree of mechanical coupling between the endoderm and adjacent paraxial mesoderm. To illustrate the broader utility of this method, we determined that forces on the order of 10 N are sufficient to unzip the neural tube during primary neurulation. Together, these findings provide basic insights into the mechanics of embryonic epithelia in vivo in the early avian embryo, and provide a useful tool for future investigations of how morphogenesis is influenced by mechanical factors.

**Graphical Abstract:** *Summary Statement:* 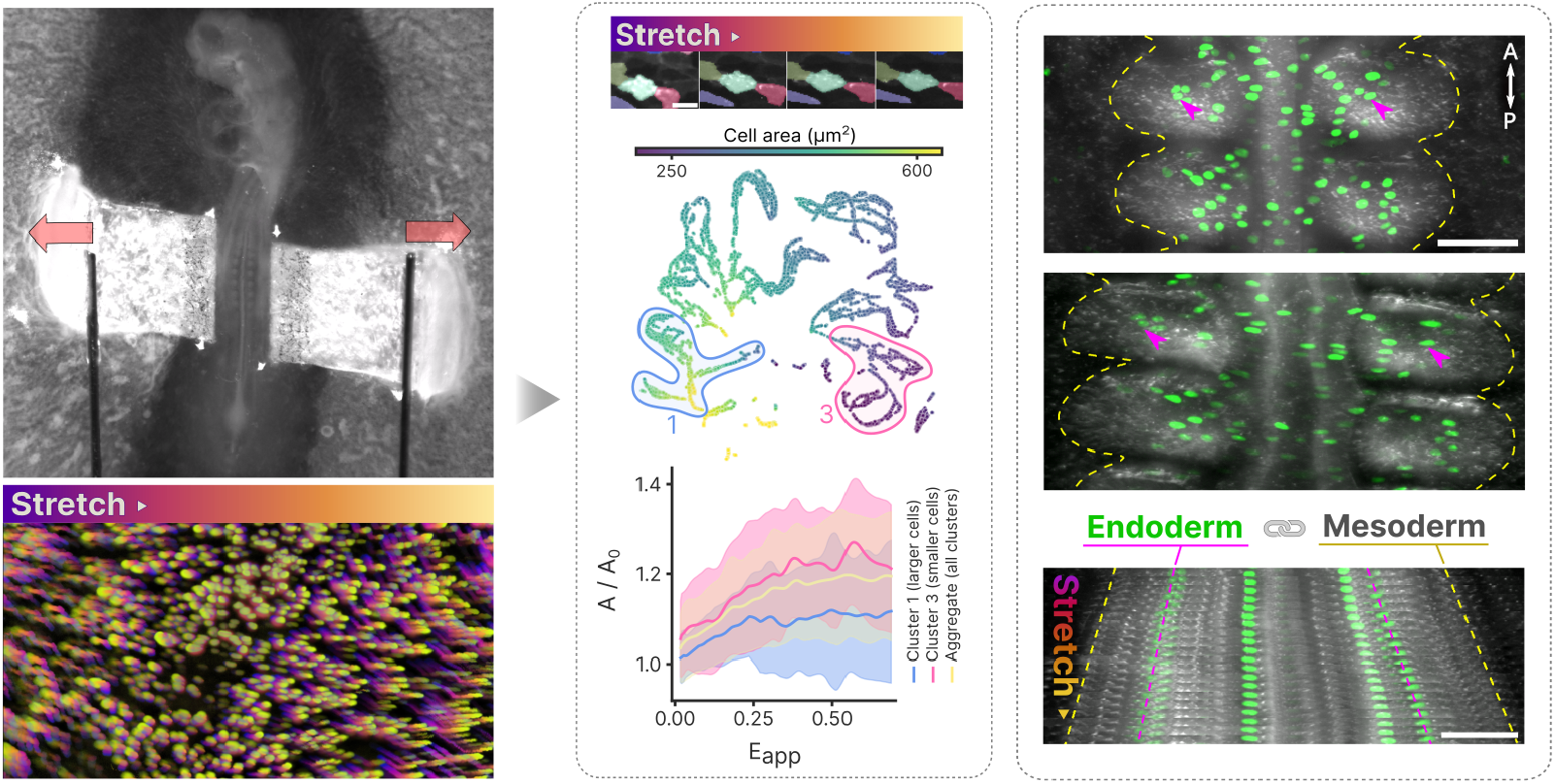 A simple approach is devised to quantitatively apply tension to epithelia in vivo, and used to study endoderm mechanics in the deeloping chick embryo.

## INTRODUCTION

Development requires coordination of cell fate specification with growth and mechanical forces that deform embryonic tissues into precise shapes. Accordingly, a mechanically motivated framework for studying development has provided increasingly cross-disciplinary insights, particularly over the past two decades. This physically-motivated view of development has revealed that mechanisms of morphogenesis can be understood using purely mechanical descriptions, providing insights even in the absence of implicating genes or signaling pathways per se (Mongera et al., 2018; Saadaoui et al., 2020; Savin et al., 2011; Serra et al., 2023; Shyer et al., 2013; Tallinen et al., 2014). At the same time, physical forces and mechanical properties have begun to be recognized as guiding cell behaviors and gene expression as well (Barriga et al., 2018; Farge, 2003). Therefore, mechanics has increasingly become woven into our understanding of morphogenesis both as an effector and modifier of developmental programs (Collinet & Lecuit, 2021). This has led to development of a range of tools for measuring forces and mechanical properties of embryonic tissue in vivo (Campàs et al., 2014; Chan et al., 2023a, 2023b; Dzementsei et al., 2022; Gómez-González et al., 2020; Mongera et al., 2018) and in explants (Barriga et al., 2018; Savin et al., 2011; Zamir et al., 2003), (Shook et al., 2018), (Zhou et al., 2009). However, these efforts have primarily focused on mesenchymal tissues, and it remains a challenge to simultaneously apply and measure tissue-scale forces in embryonic epithelia in vivo (Doubrovinski et al., 2017). As a result, mechanical behaviors of embryonic epithelia remain under-explored, particularly among amniotes.

Here, we describe a simple method to apply exogenous, uniaxial stretch to epithelia in the chick embryo in vivo. Focusing primarily on the presumptive midgut endoderm, we conducted a series of experiments investigating mechanical heterogeneity at both tissue and cell length-scales, and quantifying mechanical coupling between endoderm and mesoderm. Finally, to illustrate broader utility of the method, we applied tension to the ectoderm, quantifying the tensile forces necessary to unzip the forming neural tube.

## RESULTS AND DISCUSSION

In vivo application and measurement of tension to intact epithelia in the chick embryo To study mechanical properties of the avian endoderm, we devised a simple approach for applying tension directly onto the flat endodermal epithelium of the presumptive chick midgut. We designed an apparatus that sits on a microscope stage, and consists of a linear actuator driving bidirectional displacement of a pair of fine tungsten cantilevers to apply tension to the embryo (Fig. 1A). Cantilever bending can be used to calculate the associated force (Fig. S1A). Cantilevers of 0.13 mm diameter and 40 mm length were used to perform force measurements in the range of 1-1,000 μN. To transmit displacement of tungsten rods to endoderm stretch, we used an approach motivated by EC culture (Chapman et al., 2001), placing small L-shaped pieces of filter paper along the lateral edges of the gut-forming endoderm of Hamburger Hamilton (HH) stage 13 embryos (Fig. 1B-D). Filter paper adhered tightly to the ventral epithelium, such that when stretch was applied via lateral displacement of the paired tungsten rods, large 2-D tissue deformations could be directly observed when combined with endoderm-specific expression of a fluorescent reporter (Nerurkar et al., 2019) (Fig. 1E). Quantification of 2-D Lagrangian strains revealed a largely homogeneous deformation field, with antero-posterior shortening (negative strains, blue) accompanying extensional (positive, red) strains along the direction of applied stretch, the medio-lateral axis (Fig. 1F, G). Strains were similar between the medial gut-forming endoderm and lateral hypoblast regions (Fig. S1B), suggesting similar mechanical properties despite disparate embryonic origins and gene expression profiles (Yasunaga et al., 2005). Alignment of the directions of principal strains with the medio-lateral and antero-posterior axes of the embryo indicate that a simple biaxial deformation was achieved throughout the endoderm, with little to no shear (Fig. 1G). The endoderm was generally quite extensible, accommodating 50-70% stretch before rupturing.

**Figure 1.**
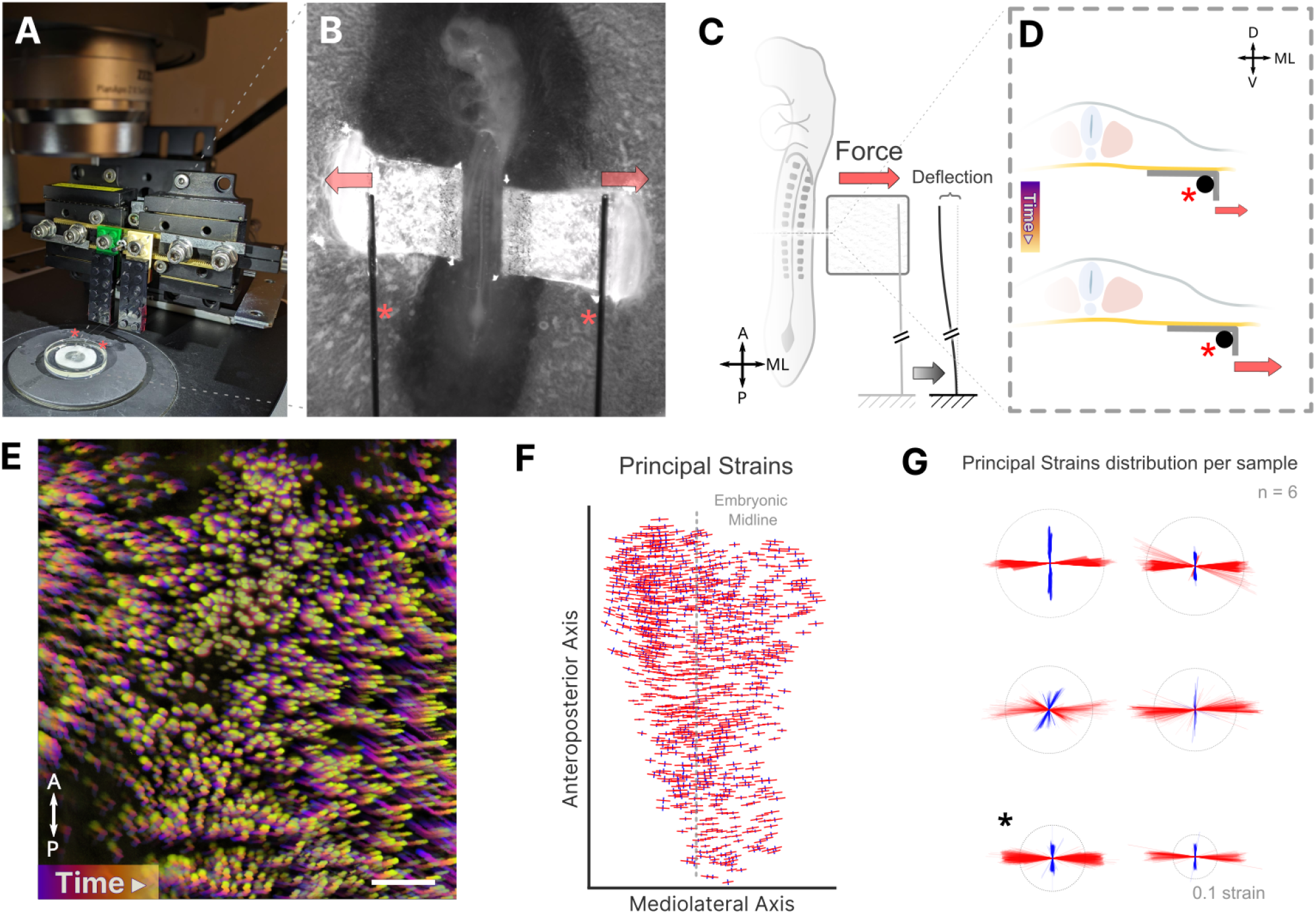
Overview of apparatus and sample stretching. (**A**) Stretching apparat us sitting atop the microscope stage, red asterisks denote the tungst encantilevers. (**B**) Magnified view of cantilevers bilaterally engaging “L-shaped” filter papers adherent to the endoderm. (**C**) Schematic representation of simultaneous stretching and force measurement via measurement of cantilever deflection. (**D**) Schematic transverse representation of the “L-shaped” filter paper attachment to endoderm and engaging the cantilever, which is displaced laterally to stretch the endoderm. (**E**) Color-coded time projection of applied deformation in endoderm following electroporation with pCAG-H2B-GFP to visualize endoderm cell nuclei; Scale = 100 *μ*m. (**F**) Principal Lagrangian strains for a representative embryo under exogenously applied medio-lateral stretch; length and orientation of red/blue lines indicat es the magnitude and orientation maximum/minimum principal strains. (**G**) Superposition of maximum (red) and minimum (blue) principal strains for 6 embryos; dashed circle indicates a magnitude of 0.1; * indicat esrepresent ative embryo from (**F**).

### Identification of single-cell mechanotypes through application of exogenous stretch

The ability to apply exogenous stretch to the endoderm in vivo provides access to a range of questions regarding the mechanical behaviors of embryonic epithelia. To begin, we asked how applied macroscopic stretch is translated locally to cell deformations, relying on endoderm-specific electroporation of pCAG-EGFP-CAAX to visualize cell boundaries during application of exogenous stretch (Fig. 2A, Supplemental Movie S1). Automated cell segmentation (Stringer et al., 2020) and tracking (Ulicna et al., 2021) was used to quantify the evolution of single-cell morphometrics for hundreds of cells per embryo as the endoderm was progressively stretched (Fig. 2B). This enables quantification of several features such as cell area, aspect ratio, orientation, strain, and position. To handle the large amount of data generated (20 features for each of approximately 20,000 cells across samples and movie frames), we employed UMAP (Uniform Manifold Approximation and Projection) to create a low-dimensional representation of high-dimensional cell morphology measurements (See Methods) (Shannon et al., 2023). Across six embryos, cells were broadly distributed in the UMAP space, reflecting the existence of distinct ‘mechanotypes’, populations of cells within the same tissue that have distinct mechanical properties (Fig. S2A). To study this cell heterogeneity further, we performed unsupervised clustering of the cells by their similarity in features (accounting for morphometry, strain, etc.) using HDBSCAN (Fig. 2C). Clusters largely included entire cell trajectories, allowing us to study the evolution of features within a cluster as a function of applied strain (Fig. S2B). Plotting each feature on UMAP space (Fig. S2C), we observed that the dominant structure in the UMAP follows an axis of increasing cell areas (Fig. 2D, Fig. S3A), with a secondary axis of separation according to applied strain and changes in cell area (Fig. S2C). For each cluster, stretching increased both cell area and aspect ratio, and led to rotation of cells toward the direction of applied stretch (Fig. S3C). However, while aspect ratio increased linearly with applied stretch, area changes were nonlinear, with cells first accommodating stretch through area expansion before plateauing to resist further area changes (Fig. 2E ; Fig. S3C). Interestingly, larger cells attenuate their area expansion at lower applied strains than smaller cells (Fig. 2E, Fig. S3B,C). This may be a consequence of the tendency for cells to preserve their volume as they deform (Wilkes & Athanasiou, 1996): cells readily shorten their apico-basal height to accommodate in-plane stretch until the balance between volumetric constraints (incompressibility of intracellular contents) and boundary constraints (cell-cell and cell-matrix contacts) begins to favor cell shape changes that do not further alter the cell height, and as a result, area.

**Figure 2.**
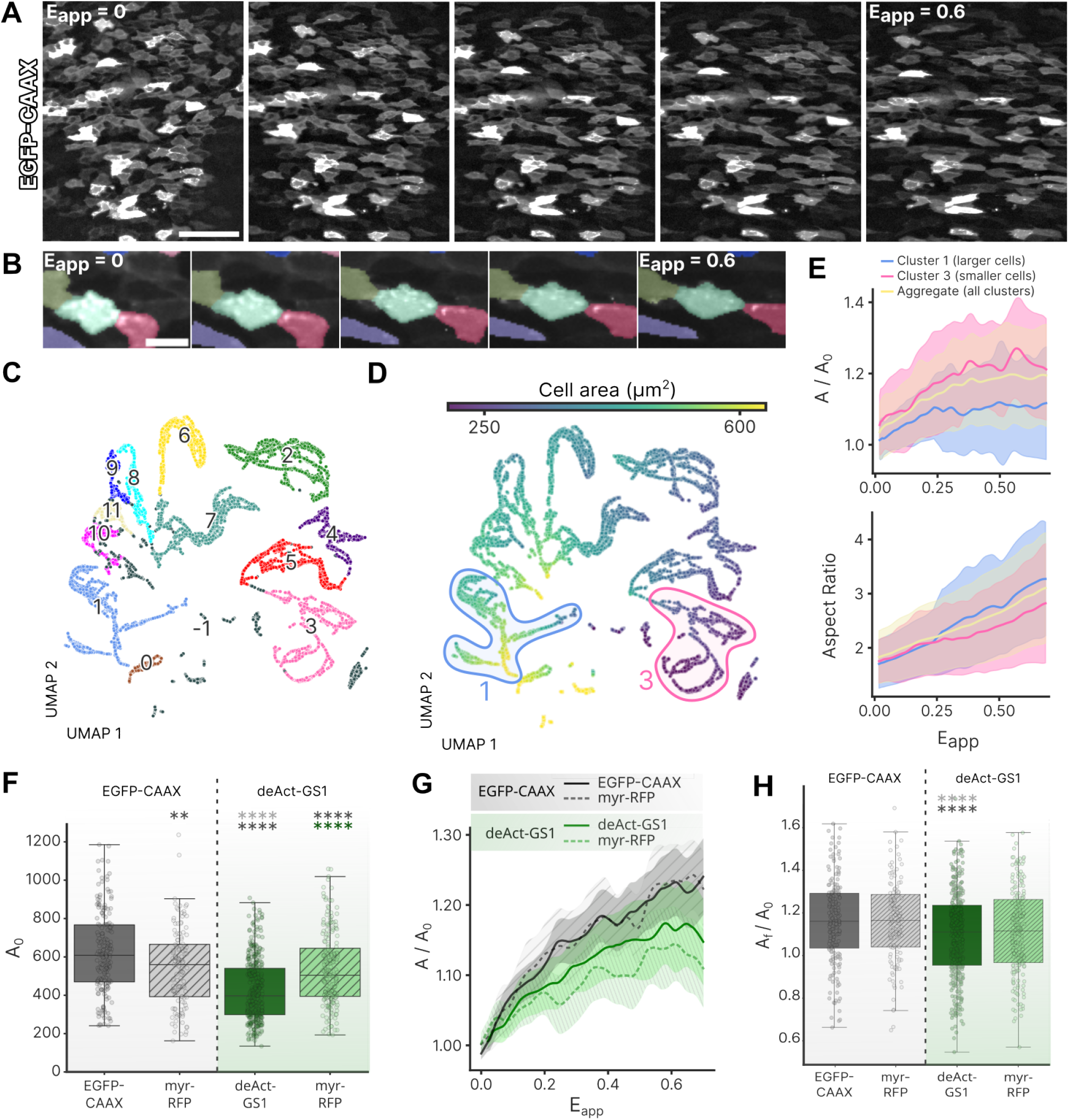
Cell scale response to applied strain reveals mechanotype heterogeneity. (**A**) Stills from a time lapse of EGFP-CAAX expressing endoderm cells as applied strain *E_app_* is increased from 0 (left) to 0.6 (right); Scale = 100 *μ*m. (**B**) Magnified view of single cell deformation as indicated by automated segmentation and tracking as *E_app_* increases from 0 (left) to 0.6 (right); Scale = 10 m (**C**) Representation of 11 distinct cell clusters identified in UMAP space; each data point indicates a single cell at a single time step during the application of exogenous stretch. (D) Cells in UMAP space color-coded according to their area. (**E**) Evolution of mean normalized cell area (*A*/*A*_0_) (top) and aspect ratio (bottom) with increasing applied strain *E_app_* for all cells, cluster 1 (large cells) and cluster 3 (small cells), and aggregate (all clusters). Solid lines indicate mean and shaded area indicates standard deviation. (**F**) Initials cell areas (*A*_0_) of EGFP-CAAX, deAct-GS1 and respective wild type (myr-RFP) neighboring cells (One-way ANOVA followed by TukeyHSD ; ** : p< 0.01 ; **** : p< 0.0001). (**G**) Evolution of mean normalized cell area (*A*/*A*_0_) with increased applied strain *E_app_* for EGFP-CAAX, deAct-GS1 and respective wild type cells. (**H**) Quantification of final increase in cell Area (*A_f_* =*A* _O_) (One-way ANOVA followed by TukeyHSD ; **** : p< 0.0001).

### Endoderm-specific disruption of F-actin attenuates cell area expansion in response to tissue-scale stretch

With the ability to apply tension to epithelia in vivo, one can test the contribution of specific subcellular components through targeted disruption of their normal function. For illustration, we focused on F-actin, as a key structural component dictating both passive and active cell properties (Bisaria et al., 2020; Brückner et al., 2019). F-actin was disrupted in a subpopulation of endoderm cells by focal electroporation to misexpress DeAct-GS1 (DeAct), a GFPfused peptide that sequesters G-actin monomers, leading to depolymerization of the F-actin (Harterink et al., 2017) (Fig. S4A). More intense disruption of F-actin using the SpvB variant of Deact (Harterink et al., 2017) resulted in delamination of cells (not shown), and we therefore focused instead on DeAct-GS1, which has a more moderate effect, for further study. To examine non cell-autonomous effects of F-actin disruption, a second electroporation was carried out to visualize neighboring cells by misexpression of a membrane localized RFP (myr-RFP) (Fig. S4B) adjacent to DeAct-GS1 expressing cells. Prior to stretching, F-actin disruption by DeAct-GS1 significantly decreased cell crosssectional area compared to neighboring myr-RFP expressing cells and compared to cells expressing membrane-localized GFP (EGFP-CAAX) in control electroporated embryos (Fig. 2F). We next examined how cell morphometrics were influenced by F-actin disruption during the application of exogenous stretch. Surprisingly, in DeAct-electroporated embryos, both myr-RFP and DeAct-GS1 expressing cells displayed limited area changes with stretch when compared to myr-RFP and EGFP-CAAX expressing cells in control embryos, respectively (Fig. 2F). This was unexpected for two reasons. First, under control conditions, cells with smaller initial area experience larger area changes under applied stretch (Fig. S3B), but smaller DeAct-GS1 cells do not follow this trend. Second, myr-RFP positive cells adjacent to DeAct-GS1 expressing cells have initial cross-sectional areas similar to control cells, but respond to stretch just as DeAct-expressing cells do, with reduced area changes as stretch is applied (Fig. 2H). These findings suggest that F-Actin may play an active role in apico-basal shortening to accommodate in-plane stretch (Sherrard et al., 2010). Interestingly, however, aspect ratio varied with macroscopic strain similarly across all cells (Fig. S4C). Making use of the ability to measure applied forces, we observed that F-actin disruption did not significantly alter relative tissue stiffness (Fig. S4D), suggesting that endoderm properties may be dominated by the stiff basement membrane rather than active cell properties. Nonetheless, these findings support an important role for F-actin in transmitting macroscopic tension in the epithelium to local cell deformations in a non cell-autonomous manner. Taken together, these findings highlight the importance of studying the mechanical response of epithelial cells in vivo, where complex interactions of cells with their neighbors and ECM may produce counter-intuitive results.

### Exogenous stretch reveals mechanical coupling between endoderm and mmesoderm

While investigating the effects of tension on in-plane properties of the endoderm, we observed, surprisingly, that as the endoderm is stretched, as much as 75% of the applied deformation was transmitted to the sub-adjacent somites (Fig. 3A-C and Supplemental Movie S3), indicating potential mechanical coupling between germ layers over large deformations and forces on the order of ∼ 100 μ N. To test whether mesodermal deformations were the result of direct mechanical coupling to the endoderm, or alternatively an indirect result of lateral fusion of dorsal and ventral extraembryonic tissues that may dorso-ventrally compress the somites as the embryo is medio-laterally stretched, we examined strain transfer between germ layers following microdissection to dorsally isolate the embryo proper from extra embryonic tissues (Fig. 3D). Following dissection, strain transfer between endoderm and somites was unaffected (Fig. 3D), suggesting that this phenomenon results directly from mechanical coupling across the interface between endoderm and somites. Similar strain transfer was observed posteriorly between the endoderm and presomitic mesoderm (Fig. S5), suggesting that mechanical coupling is more broadly present between endoderm and paraxial mesoderm as a whole (Fig. S5B).

**Figure 3.**
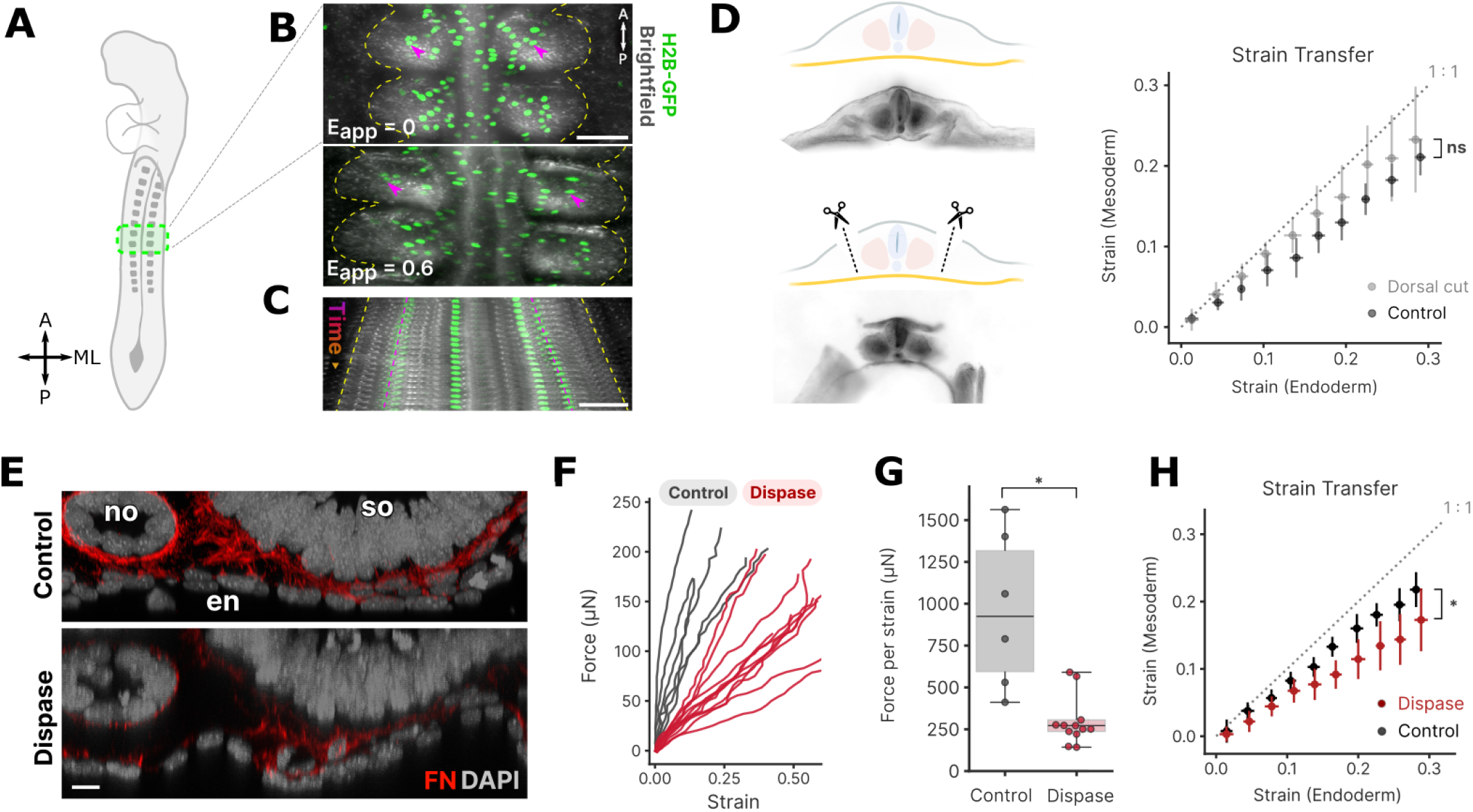
Mechanical coupling across germ layers. (A) Schematic of embryonic region imaged in (B-C), ventral view. (B) Overlay of brightfield and fluorescent images prior to (top, applied strain *E_app_* = 0) and after application of stretch to the endoderm (bottom, *E_app_* = 0.6); endoderm nuclei labeled by electroporation with pCAG-H2B-GFP (green). Magenta arrows and dashed yellow lines denote the positions of two nuclei and lateral somite boundaries, respectively; Scale = 100 *μ*m. (C) Kymograph of stretch progressively applied to endoderm, illustrating concomit ant deformation of somit es over time; Scale = 100 *μ*m. (D) Schematic (top) and brightfield (bottom) of dorsal cuts introduced to isolate ventral interactions between endoderm and somit es. At right, comparison of Lagrangian strains in endoderm and mesoderm for cont rol and dorsal cut embryos; ns = not significant (p = 0.52, Welch’s t-t est on the slopes of the strain transfer per sample). (E) Transverse view of the endoderm-somit e interface of cont rol (top) and dispase-t reated (bottom) embryos; Scale = 10 *μ*m. FN = fibronectin (red), DAPI = nuclei (grey), no = notochord, so = somite, en = endoderm. (F) Quantification of applied force vs in-plane Lagrangian strain in the endoderm for cont rol and dispase-t reated embryos. (G) Relative st iffness quant ified from force-strain curves in (F); * p = 0.012 (Welch’s t-test). (H) Comparison of Lagrangian strains in endoderm and mesoderm for cont rol and dispase-t reated embryos; * p = 0.012 (Welch’s t-t est on the slopes of the strain transfer per sample).

Based on previous identification of fibronectin pillars that span the space between the endoderm and somites (Sato et al., 2017), we next considered whether such extracellular matrix (ECM) components could be responsible for mechanical coupling. To disrupt fibronectin and collagens in the ECM, embryos were treated with dispase, (Zaher et al., 2024) significantly disrupting the fibronectin-rich ECM that forms the interface between endoderm and paraxial mesoderm (Fig. 3E). Dispase treatment produced an 80% reduction in the in-plane stiffness of the endoderm (Fig. 3F, G). Surprisingly, however, only a modest, but significant, reduction in strain transfer between endoderm and mesoderm was observed (Fig. 3H). These results suggest that fibronectin and collagen play a more dramatic role in determining the in-plane properties of the endodermal epithelium than they do in mechanical coupling between germ layers.

In light of these findings, it is somewhat surprising that exogenous stretch produces largely homogeneous strain fields throughout the midgut endoderm (Fig. 1F,G, Fig. S1B). This uniformity of endoderm deformations, despite the varied architecture and cell types of neighboring mesodermal tissues such including splanchnic, intermediate, axial, and paraxial mesoderm, is consistent with the idea that the endoderm is significantly stiffer than mesoderm. The tight mechanical coupling between PSM and a stiffer endodermal layer may have implications for a range of events such as axis elongation and gut tube morphogenesis, which have traditionally been studied as mechanically distinct events. For example, it is possible that endoderm may play a previously unappreciated role in buffering against leftright asymmetries during somitogenesis, which have been linked to scoliotic malformations during axis elongation (Pourquié, 2011).

Although mechanical interactions between germ layers have been implicated in many aspects of vertebrate morphogenesis (Smith et al., 2022), including placement of the neural anlage (Smutny et al., 2017), neurulation (Guillon et al., 2020), and axis elongation (Xiong et al., 2020), in most cases, these interactions have been inferred from mathematical modeling (Guillon et al., 2020), or indirectly by observing how tissue compartments respond to perturbations in the neighboring tissue (Smutny et al., 2017; Xiong et al., 2020). By providing access to directly assess interfacial mechanics of embryonic tissues in vivo, the present approach may therefore be of broader utility to a range of questions in morphogenesis.

### Measurement of tensile properties of the neuroepithelium during primary neurulation

To illustrate the broader utility of this method, we performed proof-of-concept experiments aimed at measuring the forces needed to unzip the neural tube during primary neurulation. Rupture of the posterior-most adhesion point in the neural tube was achieved with pulling forces on the order of ∼50 μN (Fig. 4, Supplemental Movie S4). When combined with gene misexpression and pharmacological treatments, this method could provide highly quantitative insights into the mechanobiology of neural tube zippering (Maniou et al., 2024). These results illustrate the potentially broad applications of a relatively simple, inexpensive technique for measuring tension while applying stretch to embryonic epithelia in vivo.

**Figure 4.**
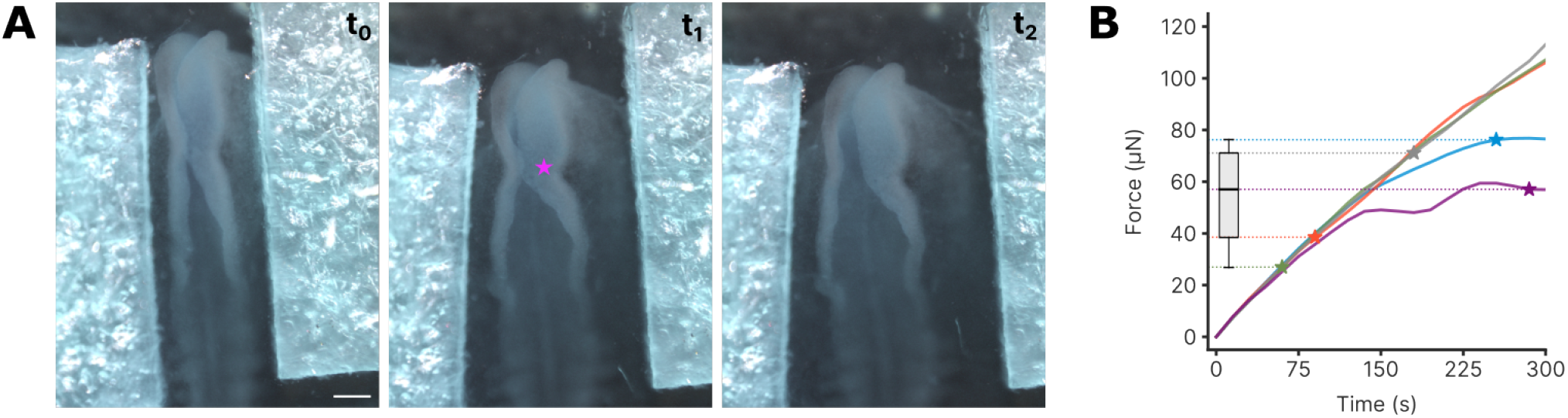
Measurement of forces necessary to reverse neural tube closure. (A) Snapshots of a HH stage 8 embryo as tension is applied across the surface ectoderm; magenta star indicates the final point of contact between the two lateral folds; Scale = 200 *μ*m. (B) Measurement of the applied force as a function of time from the start of loading; stars indicate the rupture force for a given sample. Each sample is indicated by a different color.

## MATERIALS AND M ETHODS

### Chicken embryology techniques, electroporation and drug treatments

Fertilized White Leghorn chicken (Gallus gallus domesticus) eggs were acquired from University of Connecticut Poultry Farm and incubated at 37° C and 60% humidity for 45 hours (HH stage 12). Embryos were harvested onto filter paper rings and transferred to plates containing EC culture medium (Chapman et al., 2001). Endoderm-specific electroporation was carried out as previously described (Nerurkar et al., 2019; Oikonomou et al., 2023). Briefly, electroporation solutions were prepared by diluting pCAG-H2B-EGFP, pCAG-EGFP-CAAX (Addgene # 86056), pCAG-myr-mRFP (Addgene # 32604), or pCAG-DeAct-GS1 (a fusion protein of DeAct-GS1 with EGFP, subcloned into pCAG from Addgene # 89445) plasmids to 3.5 g/L in molecular grade ddH2O with 5% sucrose and 0.1% Fast Green FCF. Following delivery of the DNA solution to the ventral surface by micropipette, embryos were electroporated using a Nepa 21 transfection system (Nepa Gene, Ichikawa City, Japan) with a sequence consisting of three 35V poring pulses of 0.2 msec duration separated by 50 msec with a decay rate of 10% between successive pulses, followed by five 4V transfer pulses of 5.0 msec duration separated by 50.0 msec with a 40% decay rate (Nerurkar et al., 2019; Oikonomou et al., 2023). Experiments were carried out after 6-8 hours incubation ex ovo, when embryos had reached HH stage 13 and fluorescent signal was readily detected. For the dorsal ectoderm dissections, embryos were placed dorsal side up under a Zeiss Stemi 508 stereo microscope. Using a flame-sharpened fine tungsten knife, bilaterally symmetric incisions were made to sever the surface ectoderm at the level of the posterior-most 4-5 somites. Embryos were then imaged as described below. To quantify strain in the PSM, tissues were labeled by microinjection of the lipophilic dye DiI (2.5 mg/mL in DMF, Invitrogen) using pulled glass capillary needles (Fig. S5A). For dispase treatment, embryos were fitted with a small plastic confining ring to retain dispase solution, and treated with 100*μ*L of a 1 U/ml dispase (Dispase I protease, Sigma-Aldrich, D4818-2MG) diluted in PBS; vehicle controls similarly received 100*μ*l of PBS. Embryos were incubated for 10 minutes at room temperature, then rinsed in PBS, and excess moisture was wicked using a Kimwipe prior to stretching experiments. Mild conditions were used to sever the ECM without completely stripping it from the tissue, as more complete dispase digestion results in severe delamination of the chick embryo (Danesin et al., 2021)

### Mechanical testing apparatus

Mechanical properties of embryonic tissues were probed using a custom micromechanical apparatus to perform uniaxial, bidirectional tensile tests. The device relies on thin tungsten rods to transmit forces to the embryo, enabling both the application of tissue strain and the measurement of the applied force (which is proportional to rod deflection, Fig. 1C, S1A). A gear system was used to achieve opposing/bidirectional displacement of a pair of tungsten rods from a single linear actuator. Each tungsten rod was afixed to a spring-loaded linear stage (Newport # 460A-X), and the two stages were in turn mounted with opposing orientations on a base plate (Newport # M-BK-3A-T). One stage was directly controlled using a linear actuator (Newport # CONEX-TRA25CC), and this movement was coupled to opposing movement of the second stage using a gear system in which a centrally-positioned gear (McMaster-Carr # 7880K14) rotates freely on a perpendicular rotary shaft (McMaster-Carr # 1327K93) to convert rightward motion from a gear rack (McMaster-Carr # 7854K11) mounted on the actuated stage into leftward motion on another gear rack that is fastened to the passive stage. On each stage, a tungsten rod of 0.127 mm diameter (A-M Systems # 716000) was attached to act as the cantilever arm. Finally, stretching of the endoderm was achieved by movement of the tungsten cantilevers against filter paper squares (2 × 2 mm) placed directly on the ventral surface of the embryo; similar experiments on the neural tube were conducted by dissecting away the vitelline membrane and placing filter paper squares on the surface ectoderm to the left and right of the forming neural tube.

Bending stiffness of the tungsten cantilever was calibrated by measuring vertical deflection of the cantilever tip following hanging small pieces of aluminium foil of with known weights from the cantilever tip. Bending stiffness (0:167 N/m) was then quantified as the slope of the force vs deflection data, as quantified by a linear regression (Fig. S1A). The tungsten cantilevers were displaced slowly at a rate of 0.003 mm/s to mitigate viscous effects. Live imaging of samples was performed as stretch was progressively applied using a ZEISS AxioZoom v16, at 50X magnification to visualize displacement of cell nuclei, and at 100X to visualize cell shape changes. Images were captured every 15-20 seconds for a total of 10-15 minutes or until tissue rupture. Imposed deformations were sufficiently large to warrant use of Lagrangian strains to quantify the deformation field, as opposed to linear/engineering strains. Accordingly, the tissue-scale applied strain was quantified as 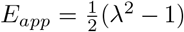, where λ = *L*/*L*_0_ is the stretch ratio relating the current/deformed distance between fiducial markers near the two filter paper squares (e.g. labelled cells) normalized to their distance prior to the onset of stretch. To report the local deformations within the epithelial plane, (Fig. 1F), the full Lagrangian strain tensor was computed as 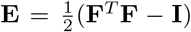, where 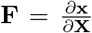 is the deformation gradient tensor and x and X are position vectors in the deformed and undeformed configuration, respectively. **F**, and subsequently **E**, were computed following the approach of (Mowlavi et al., 2022), using as input the trajectories of all cells tracked in the tissue with btrack (Ulicna et al., 2021). Direction and magnitude of principal strains were calculated by eigen-decomposition of the 2D strain tensor.

### Immunofluorescence

Embryos were fixed overnight in 4% paraformaldehyde in PBS, rinsed, and dissected from surrounding extraembryonic tissue and filter paper. Embryos were washed (PBS with 0.1% Triton was used for all washes) and left in blocking solution (10% heat inactivated goat serum) for 2 hours, then incubated overnight at 4° C with mouse anti-fibronectin primary antibody (DSHB, B3/D6) diluted at 1:200. Samples were washed and then incubated with a goat anti-mouse 488 secondary antibody (1:200) and DAPI (1:1000) overnight. Embryos were then cleared in RapiClear (RapiClear 1.49, Sunjin Labs), and imaged on a ZEISS LSM880 confocal microscope with a 40X water immersion objective, and a dense z-stack at 0.2 μm spacing for optical section reconstruction.

### Image analysis and Data processing

The code to follow the analysis pipeline is available on Github. Image analyses on time series movies obtained from the mechanical testing experiments were performed using the open-source image visualization Python library Napari (Sofroniew et al., 2022), preprocessing functions from the sci-kit image library (van der Walt et al., 2014), the segmentation algorithm Cellpose (Stringer et al., 2020) (starting from weights of the cyto2 model, trained with manual annotations for 1000 epochs on a workstation equipped with an NVIDIA T400 GPU), and btrack, a Python library for multi-object Bayesian tracking (Ulicna et al., 2021) (particle configuration). Where necessary, time lapse videos were stabilised against drift using custom code as in (Oikonomou et al., 2023), inspired by Fast4Dreg (Pylvänäinen et al., 2023). Cell morphometry features generated through this pipeline were aggregated across biological replicates and plotted in UMAP space, generally following the approach of cellPLATO (Shannon et al., 2023) and adapting code from ColabTracks (Jacquemet, 2024). Specifically, UMAP embedding was based on the following features: magnitude of applied strain, deformed aspect ratio, undeformed aspect ratio, current cell area, undeformed cell area, orientation of the cell major axis relative to the medio-lateral embryonic axis, minor axis length, major axis length, cell perimeter, and shape solidity (area / convex area). Clustering was performed on the UMAP-transformed data using HDBSCAN (McInnes et al., 2017).

### Footnotes

## Acknowledgments

We thank members of the Nerurkar lab for their valuable scientific input. We also thank Joseph Viola for help with designing and prototyping the mechanical stretching apparatus, and the image.sc forum users for their advice on image analysis tasks.

## Author Contributions

Conceptualization: P.O., L.C., N.L.N.; Methodology: P.O., L.C., H.C., N.L.N.; Formal analysis: P.O., L.C., A.E.W. H.C., N.L.N.; Investigation: P.O., L.C., H.C., A.E.W., J.F.D., N.L.N.; Resources: O.P., K.K., N.L.N.; Data curation: P.O., L.C., A.E.W., N.L.N.; Writing – original draft: P.O., L.C., A.E.W., N.L.N.; Writing – review & editing: P.O., L.C., N.L.N.; Supervision: N.L.N.; Funding acquisition: N.L.N.

## Data and Code Availability

Code to replicate the analysis can be found on Github (in the form of Python Jupyter notebooks). Representative raw image files for stretching experiments across conditions are available on Zenodo.

## SUPPLEMENTARY INFORMATION

**Figure S1.**
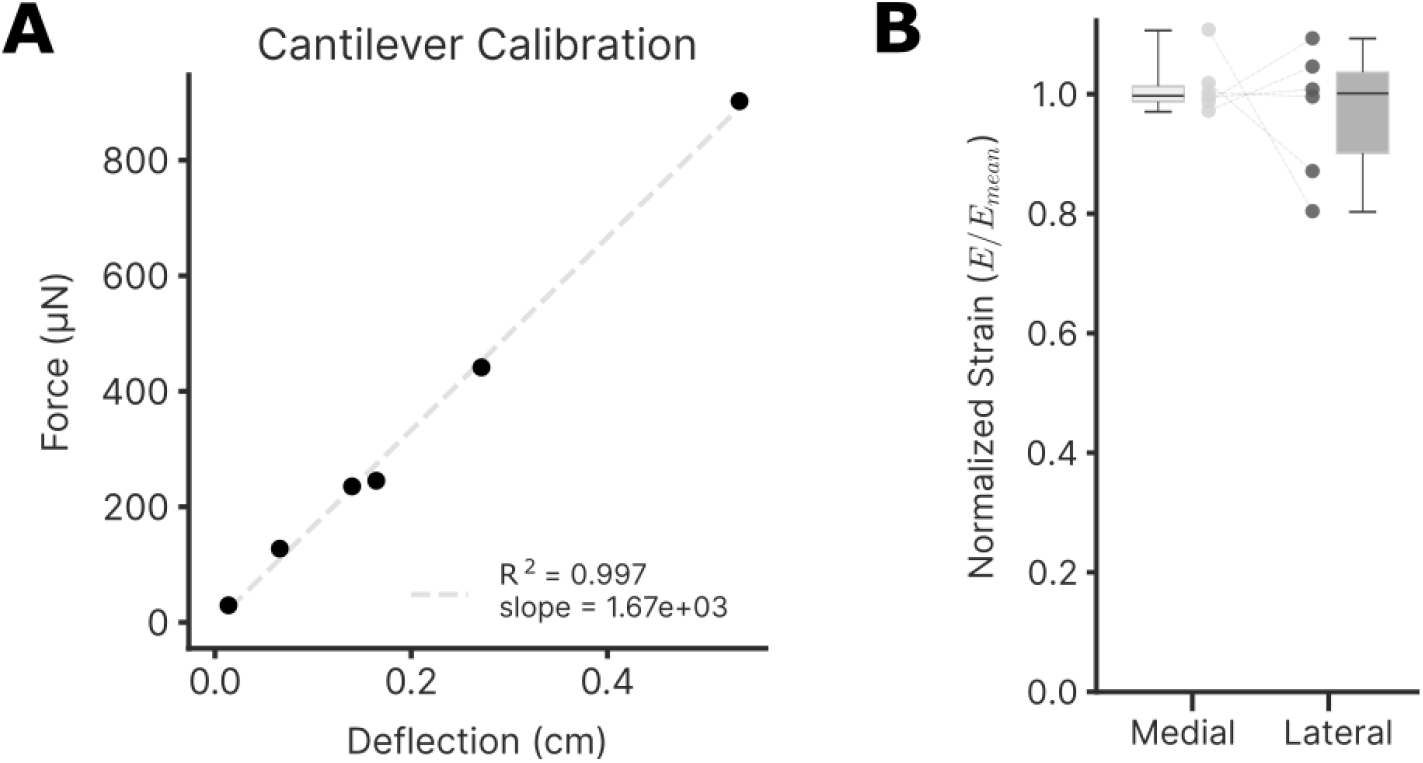
Beam calibration and strain heterogeneity. (A) Plot of cantilever deflection in function of applied force. (B) Paired comparison of mean normalized strains between medial (gut forming) and lateral endoderm in 6 different embryos.

**Figure S2.**
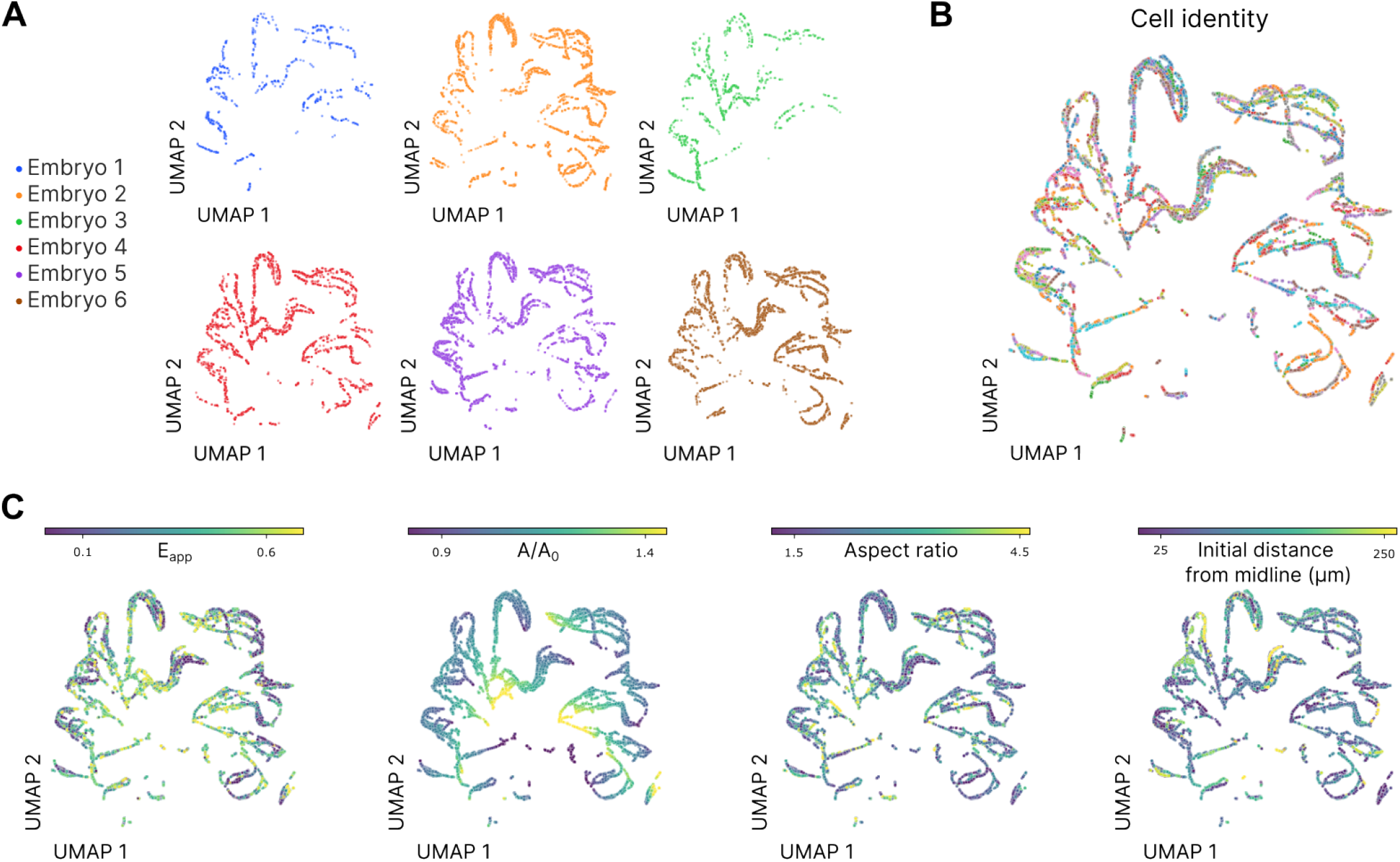
Distribution of cells in UMAP space. (A) Partitioning of cells from 6 independent experiments in UMAP space reveals each embryo distributed through the UMAP space, indicating that heterogeneity is not likely explained by embryo-to-embryo variability or batch effects. (B) Visualization of cell trajectories in UMAP space by random color assignment to individual cells throughout the course of applied deformation. (C) UMAP color-coded according to strain, cell normalized area, aspect ratio and initial distance from midline.

**Figure S3.**
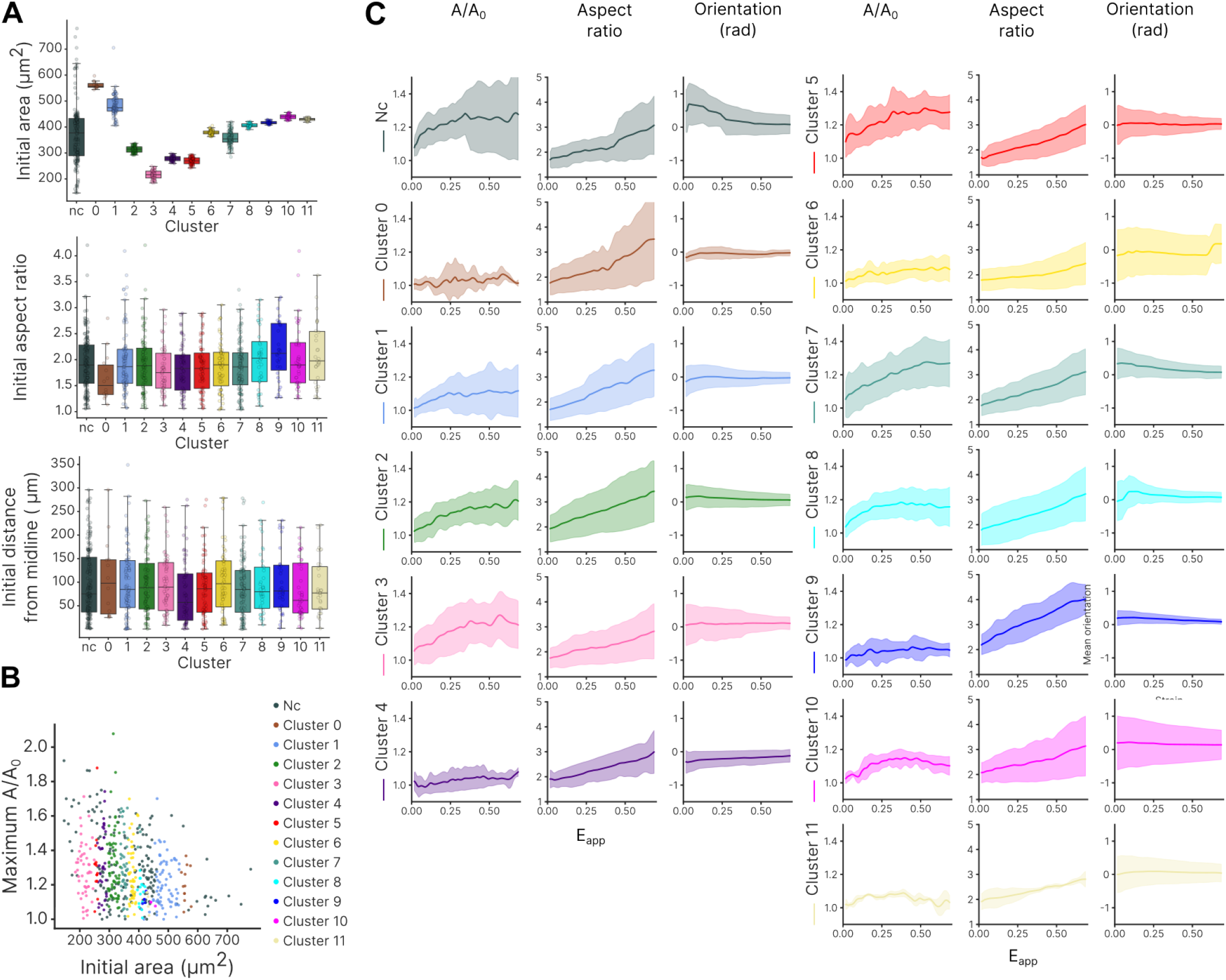
Evolution of morphometrics by mechanotype. (A) Initials features (area, aspect ratio and distance from midline) observed for the different clusters. ‘Nc’ correspond to cells that don’t belong to any cluster. (B) Maximum normalized area reached as a function of initial cell area. Colors denote corresponding cluster for each cell. (C) Normalized area, aspect ratio and orientation as a function of applied strain for each cluster. Solid lines indicate mean and shaded area indicate standard deviation.

**Figure S4.**
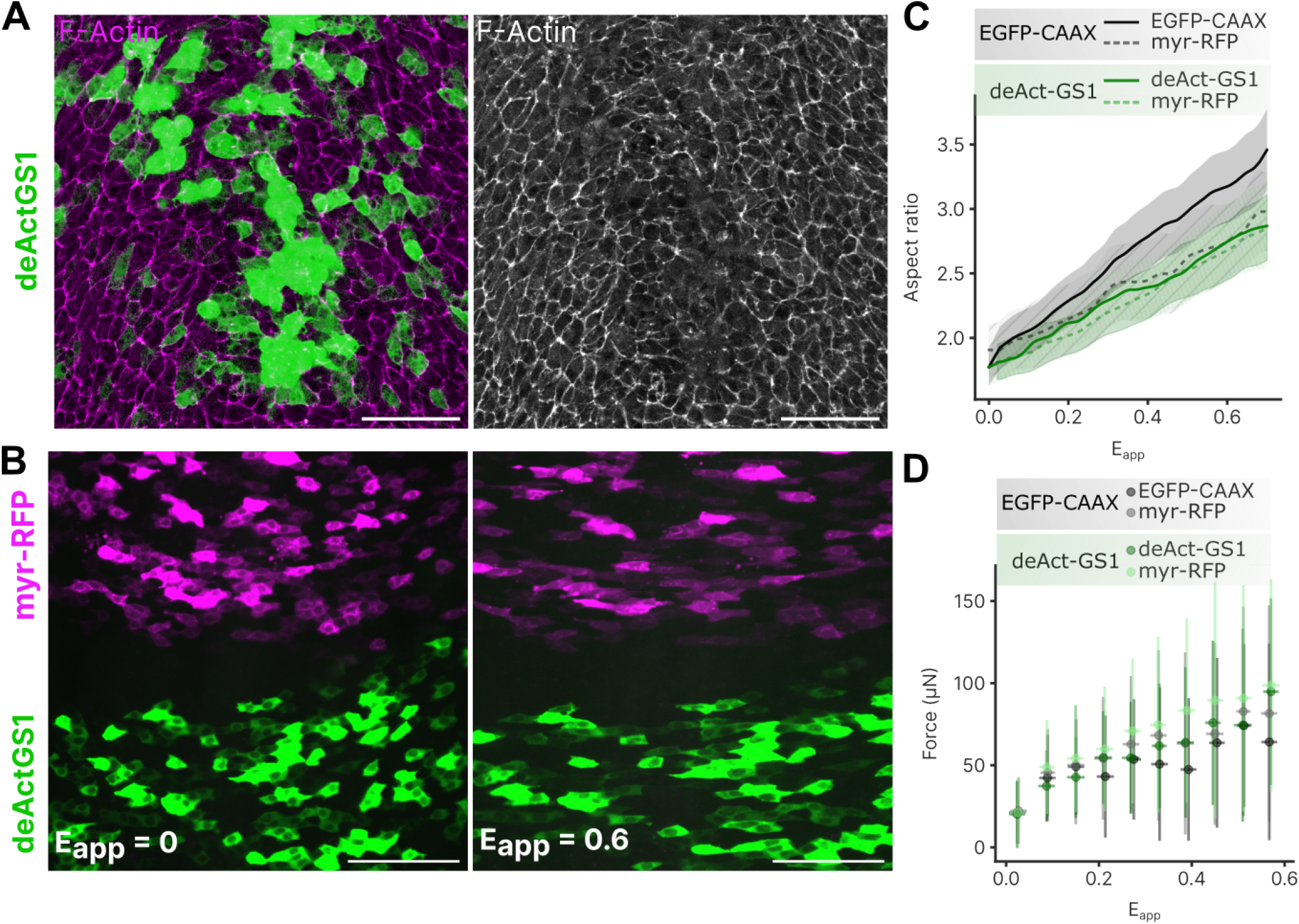
Disruption of actin through deAct-GS1 electroporation. (**A**) F-Actin staining of HH13 embryo electroporated with deAct-GS1 ; Scale = 50*µ*m. (**B**) Representative images of neighboring regions of deAct-GS1 (green) and myr-RFP (magenta) electroporated cells before stretching (left) and at 0.6 applied strain (right); Scale = 50*µ*m. (**C**) Evolution of mean cell aspect ratio with increasing applied strain *E*_*app*_ in EGFP-CAAX, deAct-GS1 and respective neighboring cells. (**D**) Applied force quantified as a function of tissue strain for control embryos electroporated with EGFP-CAAX and myr-RFP compared to embryos electroporated with deAct-GS1 and myr-RFP.

**Figure S5.**
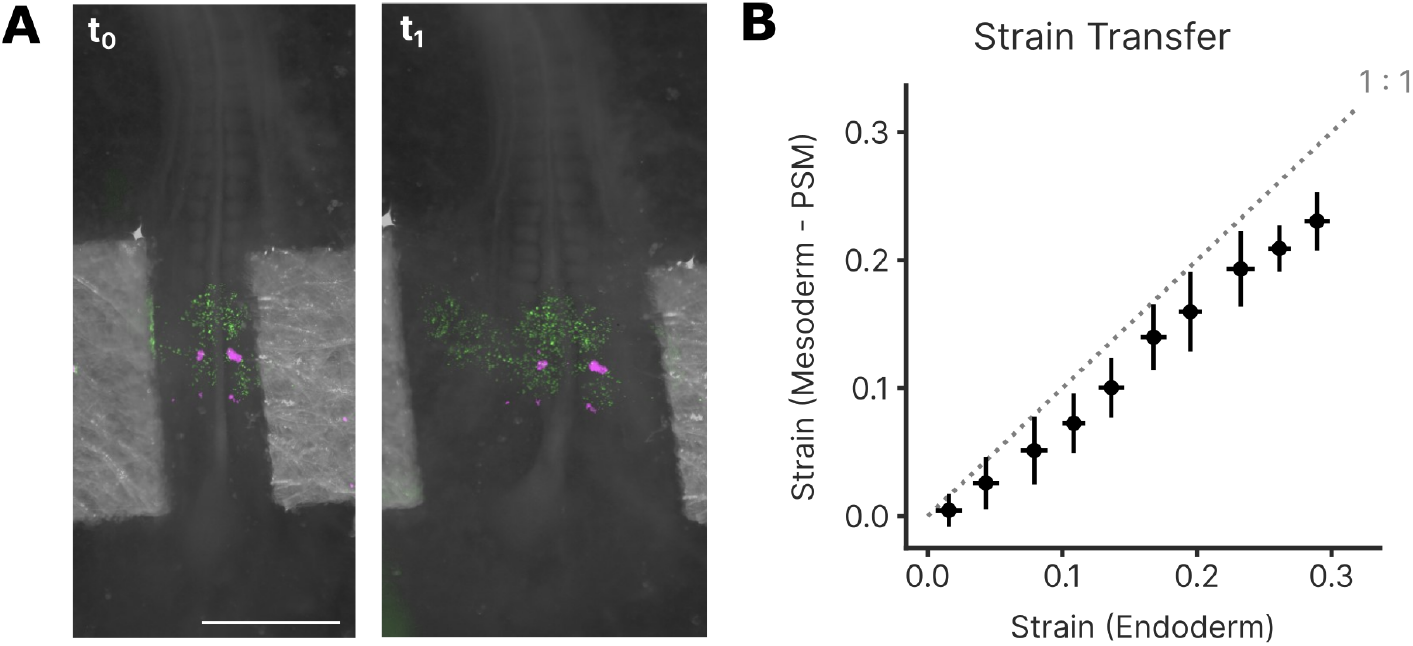
Strain transfer between endoderm and mesoderm is also observed at the PSM level. (**A**) Labelling strategy for tracking both endoderm (H2B-GFP electroporations, shown in green) and PSM (DiI injections, shown in magenta) deformations, showing pictures at the start (left) and at the end (right) of a stretching experiment. Scale = 1 mm. (**B**) Strain transfer plot between endoderm and mesoderm at the PSM level.

**Supplementary Movie 1**

`EGFP-CAAX.gif`

EGFP-CAAX stretch.

**Supplementary Movie 2**

`DeAct.gif`

DeAct stretch.

**Supplementary Movie 3**

`strain_transfer.gif`

Strain transfer.

**Supplementary Movie 4**

`neural_tube.gif`

Neural tube stretch.

## References

Barriga, E. H., Franze, K., Charras, G., & Mayor, R. (2018). Tissue stiffening coordinates morphogenesis by triggering collective cell migration in vivo. Nature, 554 (7693), 523–527. 10.1038/nature25742

Bisaria, A., Hayer, A., Garbett, D., Cohen, D., & Meyer, T. (2020). Membrane-proximal f-actin restricts local membrane protrusions and directs cell migration. Science, 368 (6496), 1205–1210. 10.1126/science.aay7794

Brückner, B. R., Nöding, H., Skamrahl, M., & Janshoff, A. (2019). Mechanical and morphological response of confluent epithelial cell layers to reinforcement and dissolution of the f-actin cytoskeleton [Physics meets medicine - at the heart of active matter]. Progress in Biophysics and Molecular Biology, 144, 77–90. 10.1016/j.pbiomolbio.2018.08.010

Campàs, O., Mammoto, T., Hasso, S., Sperling, R. A., O’Connell, D., Bischof, A. G., Maas, R., Weitz, D. A., Mahadevan, L., & Ingber, D. E. (2014). Quantifying cell-generated mechanical forces within living embryonic tissues [Publisher: Nature Publishing Group]. Nature Methods, 11 (2), 183–189. 10.1038/nmeth.2761

Chan, C. U., Xiong, F., Michaut, A., Vidigueira, J. M. N., Pourquié, O., & Mahadevan, L. (2023a). Direct force measurement and loading on developing tissues in intact avian embryos. Development, 150 (9), dev201054. 10.1242/dev.201054

Chan, C. U., Xiong, F., Michaut, A., Vidigueira, J. M. N., Pourquié, O., & Mahadevan, L. (2023b). Direct force measurement and loading on developing tissues in intact avian embryos. Development, 150 (9), dev201054. 10.1242/dev.201054

Chapman, S. C., Collignon, J., Schoenwolf, G. C., & Lumsden, A. (2001). Improved method for chick whole-embryo culture using a filter paper carrier. Developmental Dynamics: An Official Publication of the American Association of Anatomists, 220 (3), 284–289. 10.1002/1097-0177(20010301)220:3<284::AID-DVDY1102>3.0.CO;2-5

Collinet, C., & Lecuit, T. (2021). Programmed and self-organized flow of information during morphogenesis [Publisher: Nature Publishing Group]. Nature Reviews Molecular Cell Biology, 22 (4), 245–265. 10.1038/s41580-020-00318-6

Danesin, C., Ferreira, M. A., Degond, P., & Theveneau, E. (2021). Anteroposterior elongation of the chicken anterior trunk neural tube is hindered by interaction with its surrounding tissues. Cells & Development, 168, 203723. 10.1016/j.cdev.2021.203723

Doubrovinski, K., Swan, M., Polyakov, O., & Wieschaus, E. F. (2017). Measurement of cortical elasticity in drosophila melanogaster embryos using ferrofluids [Publisher: Proceedings of the National Academy of Sciences]. Proceedings of the National Academy of Sciences, 114 (5), 1051–1056. 10.1073/pnas.1616659114

Dzementsei, A., Barooji, Y. F., Ober, E. A., & Oddershede, L. B. (2022). Foregut organ progenitors and their niche display distinct viscoelastic properties in vivo during early morphogenesis stages. Communications Biology, 5 (1), 402. 10.1038/s42003-022-03349-1

Farge, E. (2003). Mechanical induction of twist in the drosophila foregut/stomodeal primordium. Current biology: CB, 13 (16), 1365–1377. 10.16/s0960-9822(03)00576-1

Gómez-González, M., Latorre, E., Arroyo, M., & Trepat, X. (2020). Measuring mechanical stress in living tissues [Publisher: Nature Publishing Group]. Nature Reviews Physics, 2 (6), 300–317. 10.1038/s42254-020-0184-6

Guillon, E., Das, D., Jülich, D., Hassan, A.-R., Geller, H., & Holley, S. (2020). Fibronectin is a smart adhesive that both influences and responds to the mechanics of early spinal column development (D. Y. Stainier, T. E. Saunders, T. E. Saunders, & J. Schwarzbauer, Eds.) [Publisher: eLife Sciences Publications, Ltd]. eLife, 9, e48964. 10.7554/eLife.48964

Harterink, M., da Silva, M. E., Will, L., Turan, J., Ibrahim, A., Lang, A. E., van Battum, E. Y., Pasterkamp, R. J., Kapitein, L. C., Kudryashov, D., Barres, B. A., Hoogenraad, C. C., & Zuchero, J. B. (2017). DeActs: Genetically encoded tools for perturbing the actin cytoskeleton in single cells. Nature methods, 14 (5), 479–482. 10.1038/nmeth.4257

Jacquemet, G. (2024, January 22). CellTracksColab—a platform for compiling, analyzing, and exploring tracking data [Pages: 2023.10.20.563252 Section: New Results]. 10.1101/2023.10.20.563252

Maniou, E., Todros, S., Urciuolo, A., Moulding, D. A., Magnussen, M., Ampartzidis, I., Brandolino, L., Bellet, P., Giomo, M., Pavan, P. G., Galea, G. L., & Elvassore, N. (2024). Quantifying mechanical forces during vertebrate morphogenesis. Nature Materials, 23 (11), 1575–1581. 10.1038/s41563-024-01942-9

McInnes, L., Healy, J., & Astels, S. (2017). Hdbscan: Hier-archical density based clustering. Journal of Open Source Software, 2 (11), 205. 10.21105/joss.00205

Mongera, A., Rowghanian, P., Gustafson, H. J., Shelton, E., Kealhofer, D. A., Carn, E. K., Serwane, F., Lucio, A. A., Giammona, J., & Campàs, O. (2018). A fluid-to-solid jamming transition underlies vertebrate body axis elongation [Publisher: Nature Publishing Group]. Nature, 561 (7723), 401–405. 10.1038/s41586-018-0479-2

Mowlavi, S., Serra, M., Maiorino, E., & Mahadevan, L. (2022). Detecting lagrangian coherent structures from sparse and noisy trajectory data. Journal of Fluid Mechanics, 948, A4. 10.1017/jfm.2022.652

Nerurkar, N. L., Lee, C., Mahadevan, L., & Tabin, C. J. (2019). Molecular control of macroscopic forces drives formation of the vertebrate hindgut [Number: 7740 Publisher: Nature Publishing Group]. Nature, 565 (7740), 480–484. 10.1038/s41586-018-0865-9

Oikonomou, P., Cirne, H. C., & Nerurkar, N. L. (2023, May 18). A chemo-mechanical model of endoderm movements driving elongation of the amniote hindgut [Pages: 2023.05.18.541363 Section: New Results]. 10.1101/2023.05.18.541363

Pourquié, O. (2011). Vertebrate segmentation: From cyclic gene networks to scoliosis. Cell, 145 (5), 650–663. 10.1016/j.cell.2011.05.011

Pylvänäinen, J. W., Laine, R. F., Saraiva, B. M. S., Ghimire, S., Follain, G., Henriques, R., & Jacquemet, G. (2023). Fast4dreg - fast registration of 4d microscopy datasets. Journal of Cell Science, 136 (4). 10.1242/jcs.260728

Saadaoui, M., Rocancourt, D., Roussel, J., Corson, F., & Gros, J. (2020). A tensile ring drives tissue flows to shape the gastrulating amniote embryo [Publisher: American Association for the Advancement of Science]. Science, 367 (6476), 453–458. 10.1126/science.aaw1965

Sato, Y., Nagatoshi, K., Hamano, A., Imamura, Y., Huss, D., Uchida, S., & Lansford, R. (2017). Basal filopodia and vascular mechanical stress organize fibronectin into pillars bridging the mesoderm-endoderm gap. Development, 144 (2), 281–291. 10.1242/dev.141259

Savin, T., Kurpios, N. A., Shyer, A. E., Florescu, P., Liang, H., Mahadevan, L., & Tabin, C. J. (2011). On the growth and form of the gut [Publisher: Nature Publishing Group]. Nature, 476 (7358), 57–62. 10.1038/nature10277

Serra, M., Serrano Nájera, G., Chuai, M., Plum, A. M., Santhosh, S., Spandan, V., Weijer, C. J., & Mahadevan, L. (2023). A mechanochemical model recapitulates distinct vertebrate gastrulation modes [Publisher: American Association for the Advancement of Science]. Science Advances, 9 (49), eadh8152. 10.1126/sciadv.adh8152

Shannon, M. J., Eisman, S. E., Lowe, A. R., Sloan, T., & Mace, E. M. (2023, October 29). cellPLATO: An unsupervised method for identifying cell behaviour in heterogeneous cell trajectory data (preprint). Cell Biology. 10.1101/2023.10.28.564355

Sherrard, K., Robin, F., Lemaire, P., & Munro, E. (2010). Sequential activation of apical and basolateral contractility drives ascidian endoderm invagination. Current biology: CB, 20 (17), 1499–1510. 10.1016/j.cub.2010.06.075

Shook, D. R., Kasprowicz, E. M., Davidson, L. A., & Keller, R. (2018). Large, long range tensile forces drive convergence during xenopus blastopore closure and body axis elongation (T. C. McDevitt, Ed.) [Publisher: eLife Sciences Publications, Ltd]. eLife, 7, e26944. 10.7554/eLife.26944

Shyer, A. E., Tallinen, T., Nerurkar, N. L., Wei, Z., Gil, E. S., Kaplan, D. L., Tabin, C. J., & Mahadevan, L. (2013). Villification: How the gut gets its villi. Science, 342 (6155), 212–218. 10.1126/science.1238842

Smith, S. J., Guillon, E., & Holley, S. A. (2022). The roles of inter-tissue adhesion in development and morphological evolution. Journal of Cell Science, 135 (9), jcs259579. 10.1242/jcs.259579

Smutny, M., Ákos, Z., Grigolon, S., Shamipour, S., Ruprecht, V., Čapek, D., Behrndt, M., Papusheva, E., Tada, M., Hof, B., Vicsek, T., Salbreux, G., & Heisenberg, C.-P. (2017). Friction forces position the neural anlage. Nature Cell Biology, 19 (4), 306–317. 10.1038/ncb3492

Sofroniew, N., Lambert, T., Evans, K., Nunez-Iglesias, J., Bokota, G., Winston, P., Peña-Castellanos, G., Yamauchi, K., Bussonnier, M., Doncila Pop, D., Can Solak, A., Liu, Z., Wadhwa, P., Burt, A., Buckley, G., Sweet, A., Migas, L., Hilsenstein, V., Gaifas, L., … McGovern, A. (2022, May 31). Napari: A multi-dimensional image viewer for python (Version v0.4.16). Zenodo. 10.5281/zenodo.6598542

Stringer, C., Wang, T., Michaelos, M., & Pachitariu, M. (2020, February 3). Cellpose: A generalist algorithm for cellular segmentation (preprint). Bioinformatics. 10.1101/2020.02.02.931238

Tallinen, T., Chung, J. Y., Biggins, J. S., & Mahadevan, L. (2014). Gyrification from constrained cortical expansion. Proceedings of the National Academy of Sciences, 111 (35), 12667–12672. 10.1073/pnas.1406015111

Ulicna, K., Vallardi, G., Charras, G., & Lowe, A. R. (2021). Automated deep lineage tree analysis using a bayesian single cell tracking approach [Publisher: Frontiers]. Frontiers in Computer Science, 3. 10.3389/fcomp.2021.734559

van der Walt, S., Schönberger, J. L., Nunez-Iglesias, J., Boulogne, F., Warner, J. D., Yager, N., Gouillart, E., Yu, T., & contributors the scikit-image, t. s.-i. (2014). Scikit-image: Image processing in python [Publisher: arXiv Version Number: 1]. 10.48550/ARXIV.1407.6245

Wilkes, R. P., & Athanasiou, K. A. (1996). The intrinsic incompressibility of osteoblast-like cells. Tissue Engineering, 2 (3), 167–181. 10.1089/ten.1996.2.167

Xiong, F., Ma, W., Bénazéraf, B., Mahadevan, L., & Pourquié, O. (2020). Mechanical coupling coordinates the co-elongation of axial and paraxial tissues in avian embryos. Developmental Cell, 55 (3), 354–366.e5. 10.1016/j.devcel.2020.08.007

Yasunaga, M., Tada, S., Torikai-Nishikawa, S., Nakano, Y., Okada, M., Jakt, L. M., Nishikawa, S., Chiba, T., Era, T., & Nishikawa, S.-I. (2005). Induction and monitoring of definitive and visceral endoderm differentiation of mouse ES cells. Nature Biotechnology, 23 (12), 1542–1550. 10.1038/nbt1167

Zaher, M., Yelin, R., Arraf, A. A., Jadon, J., Asleh, M. A., Goltzman, S., Shaulov, L., Reinhardt, D. P., & Schultheiss, T. M. (2024, April). Stored Elastic Bending Tension as a Mediator of Embryonic Body Folding. 10.1101/2024.04.02.587781

Zamir, E. A., Srinivasan, V., Perucchio, R., & Taber, L. A. (2003). Mechanical asymmetry in the embryonic chick heart during looping. Annals of Biomedical Engineering, 31 (11), 1327–1336. 10.1114/1.1623487

Zhou, J., Kim, H. Y., & Davidson, L. A. (2009). Actomyosin stiffens the vertebrate embryo during crucial stages of elongation and neural tube closure. Development (Cambridge, England), 136 (4), 677–688. 10.1242/dev.026211

